# Single-domain antibody delivery using an mRNA platform protects against lethal doses of botulinum neurotoxin A

**DOI:** 10.1101/2022.10.29.514343

**Authors:** Eugenia A. Panova, Denis A. Kleymenov, Dmitry V. Shcheblyakov, Evgeniia N. Bykonia, Elena P. Mazunina, Alina S. Dzharullaeva, Anastasia N. Zolotar, Artem A. Derkaev, Ilias B. Esmagambetov, Ivan I. Sorokin, Evgeny V. Usachev, Igor A. Ivanov, Timofei S Zatsepin, Sergey E. Dmitriev, Vladimir A. Gushchin, Denis Y. Logunov, Alexander L. Gintsburg

**Affiliations:** National Research Centre for Epidemiology and Microbiology named after Honorary Academician N F Gamaleya of the Ministry of Health of the Russian Federation, 123098 Moscow, Russia; Belozersky Institute of Physico-Chemical Biology, Lomonosov Moscow State University, Moscow, 119234 Russia; Faculty of Bioengineering and Bioinformatics, Lomonosov Moscow State University, Moscow, 119234 Russia; Center for Life Sciences, Skolkovo Institute of Science and Technology, Moscow 143026, Russia; Chemistry Department, Lomonosov Moscow State University, Moscow 119992, Russia; Department of Virology, Lomonosov Moscow State University, 119991 Moscow, Russia; Infectiology Department, I. M. Sechenov First Moscow State Medical University, 119146 Moscow, Russia

## Abstract

Single-domain antibodies (sdAbs, VHHs, or nanobodies) are a promising tool for the treatment of both infectious and somatic diseases. Their small size greatly simplifies any genetic engineering manipulations. Such antibodies have the ability to bind hard-to-reach antigenic epitopes through long parts of the variable chains, the third complementarity-determining regions (CDR3s). VHH fusion with the canonical immunoglobulin Fc fragment allows the Fc-fusion single-domain antibodies (VHH-Fc) to significantly increase their neutralizing activity and serum half-life. Previously we have developed and characterized VHH-Fc specific to botulinum neurotoxin A (BoNT/A), that showed a 1000-fold higher protective activity than monomeric form when challenged with five times the lethal dose (5 LD_50_) of BoNT/A. During the COVID-19 pandemic, mRNA vaccines based on lipid nanoparticles (LNP) as a delivery system have become an important translational technology that has significantly accelerated the clinical introduction of mRNA platforms. We have developed an mRNA platform that provides long-term expression after both intramuscular and intravenous application. The platform has been extensively characterized using firefly luciferase (Fluc) as a reporter. An intramuscular administration of LNP-mRNA encoding VHH-Fc antibody made it possible to achieve its rapid expression in mice and resulted in 100% protection when challenged with up to 100 LD_50_ of BoNT/A. The presented approach for the delivery of sdAbs using mRNA technology greatly simplifies drug development for antibody therapy and can be used for emergency prophylaxis.

## INTRODUCTION

Botulism is a rare but severe and often fatal disease caused by botulinum neurotoxin (BoNT) produced by bacterium *Clostridium botulinum*. Botulism most commonly occurs following ingestion of contaminated food, penetration of spores through the wound or when spores are ingested and then germinate in the intestinal tract (more common in infants) [1]. The inhalation of the pre-formed toxin also causes disease, but this event does not happen naturally and may occur in the context of a bioterrorist attack. Botulinum toxin has a history of use as a potential bioweapon, and thus it is classified as a Category A bioterror agent [2]. Among the four major human pathogenic BoNTs (BoNT/A, B, E, and F), produced by different serotypes of *C. botulinum*, BoNT/A poses the most serious threat for human due to its high potency and long duration of action [3]. Due to the potency and rapid onset of symptoms, exposure to botulinum toxin demands immediate anti-toxin therapy, which is usually based on equine antitoxin sera. The prophylactic therapy is also required for an individual who has eaten food suspected of being infected with *C. botulinum*. However, side effects, such as allergic reactions, fever serum sickness, and anaphylactic reactions, often occur [4]. A vaccine against botulism exists but it is rarely used as its effectiveness has not been fully evaluated and it has demonstrated negative side effects [5].

An alternative to antitoxin serum with minimal or no side effects is treatment with monoclonal antibodies (mAbs) to BoNT [6,7]. Along with conventional antibodies, single-domain antibodies (sdAb), also referred to as VHHs (variable domains of heavy-chain only antibodies) or nanobodies, opened new perspectives in the engineering of antibodies since they were discovered [8]. VHHs have some advantages over classical immunoglobulins (IgG), such as simplicity of engineering, improved heat and pH stability, and better tissue penetration [9]. However, the problems of widespread use of mAbs (both classical IgG and nanobodies) for the treatment of infectious diseases are mainly associated with the complexity and high cost of producing recombinant antibody proteins (i.e. production in cultured mammalian cells followed by purification from complex media). Furthermore, these proteins have complex post-translational modifications, which can strongly impact on their biological activity and therapeutic properties of the mAbs. Thus, they need costly development and implementation of numerous analytical tools for accurate downstream control [10].

Delivery of DNA or mRNA that encode mAbs is an attractive way to circumvent the problems of the recombinant mAb production and purification [11,12]. The possibility of using mRNA-encoded antibodies may bring some advantages, such as no risk of genome integration, more controlled exposure and more rapid protein production compared to DNA-based vectors [10]. Over the past decade, mRNA-based drugs have demonstrated a high potential in the treatment of various diseases [13–15]. During the COVID-19 pandemic, the possibility of rapid development of mRNA vaccines based on lipid nanoparticles (LNP) as a delivery system and their successful implementation were demonstrated. This breakthrough made a significant contribution to limiting the spread of the virus and reducing or stabilizing the COVID-19 incidence in countries where these vaccines were used [16,17]. These recent advances in the treatment and prevention of infectious diseases using mRNA-based technology have increased the interest in antibody production based on this method for both therapeutic and prophylactic purposes [12,18,19].

Here we present an LNP-mRNA platform providing efficient long-term expression of a delivered gene *in vivo*. First, we showed a rapid and sustained accumulation of a reporter protein (firefly luciferase, Fluc) in mice through both intramuscular and intravenous administration routes. We then applied this technology for the production of a highly protective BoNT/A-specific nanobody fused to the human Fc fragment (hereafter called “B11-Fc antibody”), which we described previously [20]. We report 100% protection in mice after intramuscular administration of the LNP-mRNA preparation as early as 3.5 h before the injection of five times the lethal dose (5 LD_50_) of BoNT/A. In the case of LNP-mRNA encoding VHH-Fc application 48 hours before challenge a 100% of the animals were resistant to 100 LD_50_ dose. The described approach greatly simplifies the development and widespread use of drugs based on sdAbs and can be used for emergency prophylaxis of various toxic and infectious diseases.

## RESULTS

### Design mRNA platform and its characterization in cell cultures and living mice

To develop an mRNA platform for efficient and long-term expression, we took advantage of fleeting mRNA transfection (FLERT) of cultured mammalian cells we have successfully used before [21,22]. To provide a high translation rate and stability, all *in vitro* transcribed mRNAs had (i) the cap-1 structure at the 5′ end; (ii) a 100-nt long poly(A)-tail at the 3′ end; (iii) the 5′ and 3′ untranslated regions (UTRs) from the human hemoglobin alpha subunit (*HBA1*) mRNA; (iv) codon optimized coding sequences (CDS), as schematically shown in **Figure 1a**. A modified nucleoside, N1-methylpseudouridines (m^1^Ψ), were co-transcriptionally incorporated into the mRNA instead of 100% uridines (U) to reduce mRNA immunogenicity [23,24].

**Figure 1.**
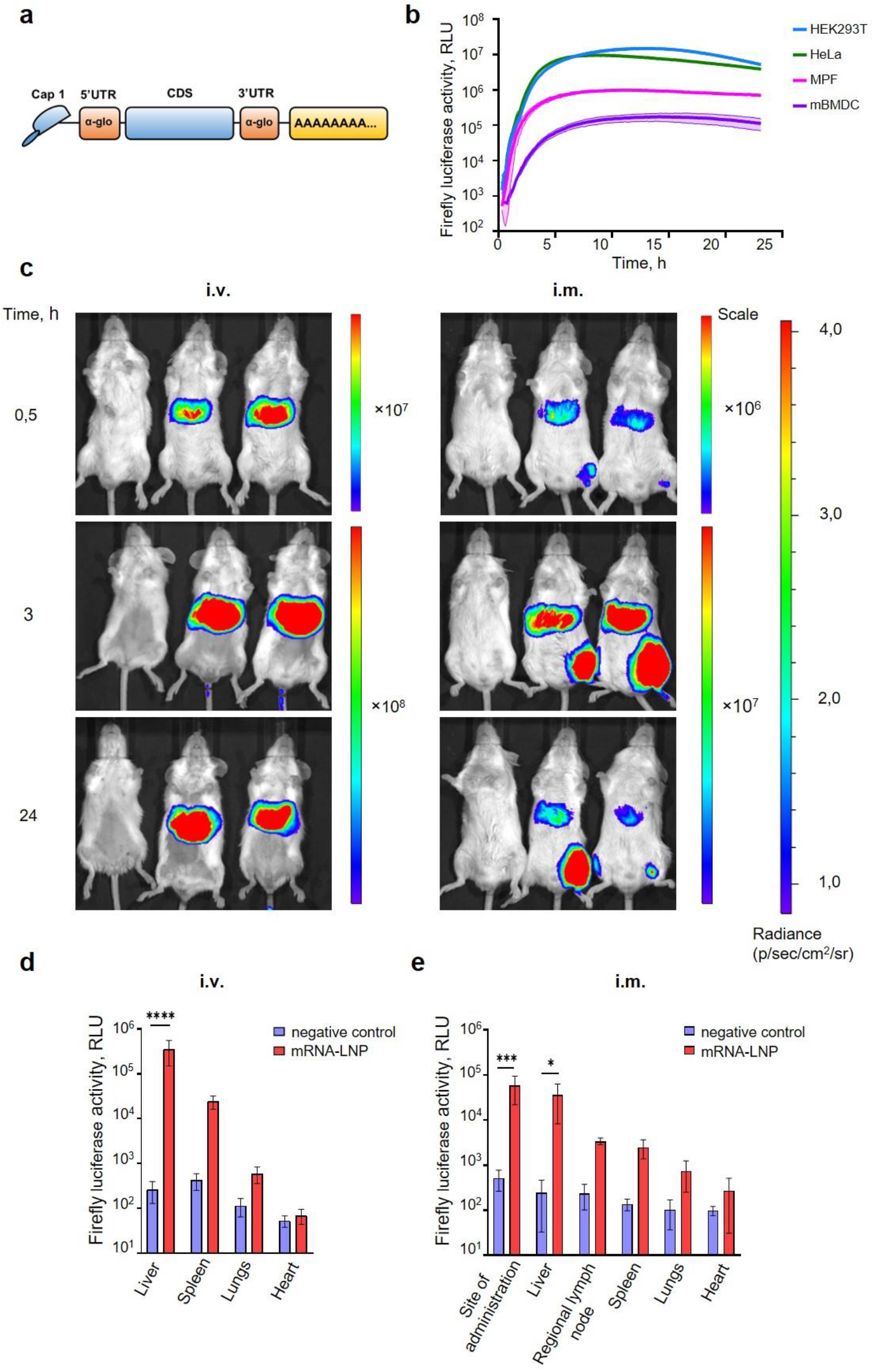
An mRNA platform design and characterization. **a**. Schematic representation of the mRNA platform used in this study. **b**. Firefly luciferase expression continuously measured after mRNA transfection of cultured mammalian cells: two human cell lines (HEK293T and HeLa), mouse primary fibroblasts (MPF), and mouse bone marrow-derived dendritic cells (mBMDC). Representative curves of at least three independent experiments are shown, expresses as mean (lines) ± SD (shaded regions) of three technical replicates from the same experiment. **c**. Luciferase activity in mice detected with *in vivo* imaging system (IVIS). BALB/c mice were injected with 5.0 μg mRNA-LNP in PBS, or PBS (for placebo group) by intramuscular (i.m.) and intravenous (i.v.) routes. Relative luminescence plot is shown, and the scale of luminescence is indicated. **d**. and **e**. Luciferase activity in homogenates of different organs after 3 h after intramuscular (i.m.) or intravenous (i.v.) administration of mRNA-LNP preparation. Boxes and whiskers represent means with standard deviations (SD). *p*-value determined by Bonferroni’s multiple comparisons test. *\*p* < 0.05, *\*\*p* < 0.01, *\*\*\*p* < 0.001, *\*\*\*\*p* < 0.0001.

First, the efficiency of the designed mRNA platform was explored with firefly luciferase (Fluc). Different cell cultures, including cancer and non-cancer human cell line (HeLa and HEK293T), as well as primary cells: mouse primary fibroblasts (MPF) and mouse bone marrow-derived dendritic cells (mBMDC), were transfected with the Fluc-encoding mRNA. Luciferase activity was continuously measured within 24 h in a temperature controlled plate reader with CO_2_ supply, as described previously [25]. In all cases, we observed a pronounced expression of the transfected mRNA, although at different levels and duration (**Figure 1b**). As expected [24,26], in primary cells (in particular, in mBMDCs) the efficient expression strictly required both 100% m^1^Ψ incorporated instead of uridine, and the presence of cap-1 instead of cap-0 (**Figure S1**).

To investigate the rate and distribution of the protein production from Fluc-mRNA, mRNA-LNP were prepared (see Methods for details) and administered into mice using two delivery routes: intravenous (tail vein) and intramuscular (posterolateral thigh muscle group) injections. The luciferase activity was detected as early as 15 min in the liver after intravenous (i.v.) mRNA-LNP injection, as well as in the site of injection for intramuscular (i.m.) route of administration (data not shown). 30 min after i.m. administration, the bioluminescence was observed mainly in the site of injection and in the liver (confirming a systemic spread of the mRNA-LNPs), but at a lower level compared to the i.v. administration (**Figure 1c**). For i.v. injection, bioluminescence has been raised within the first three hours post injection and then decreased slightly by the 24 h. For i.m. delivery route, a sharp increase in the Fluc activity was also observed during the first 3 hours post injection, after that it was declined in the liver and was not detected by the third day (**Figure S2**), whereas a very long-term expression (up to 21 days post-injection) was observed in the site of injection (**Figure S2**).

Distribution of Fluc-mRNA expression was further confirmed by assaying luciferase activity in homogenates of isolated organs (**Figure 1d, e**). The highest luciferase activity was detected in the liver homogenates for i.v. route of administration, as well as in the muscle and liver homogenates for i.m. route of injection.

### Passive B11-mRNA-LNP immunization and BoNT/A lethal dose protection

Based on the previous study of the camelid nanobodies [20], we designed mRNA coding for dimer of BoNT/A-specific camelid VHHs fused to human IgG Fc-fragment. The mRNA was assembled into LNP (B11-mRNA-LNP) according to the same protocol as for Fluc-mRNA before.

To assess the prophylactic effect of the single administration of B11-mRNA-LNP, the BoNT/A challenge experiments were performed. Three groups of mice (n = 3) were treated with B11-mRNA-LNP (10 μg per mouse), the fourth group was treated with recombinant antibody B11-Fc (10 μg per mouse), and the fifth one (the negative control) was treated with sterile PBS. Then all animals were challenged with 5 LD_50_ dose of BoNT/A according to the schedule (**Figure 2a**). Groups treated with B11-mRNA-LNP differed from each other in the time interval between prophylaxis and challenge (0, 3.5, 8 or 24 h). Mice in groups pre-treated with PBS or B11-mRNA-LNP at 0 h interval had been shown moderate symptoms in four hours and then died within 8 h and 12 h after BoNT/A administration, respectively. Group of mice with 3.5 h interval between prophylaxis and challenge showed mild clinical symptoms 26 h after intoxication but were survived up to 72 h without deterioration. Other groups of mice that received B11-mRNA-LNP or recombinant B11-Fc antibody and subsequently were challenged in 8 and 24 h, survived and displayed no symptoms of botulism (**Figure 2b**).

**Figure 2.**
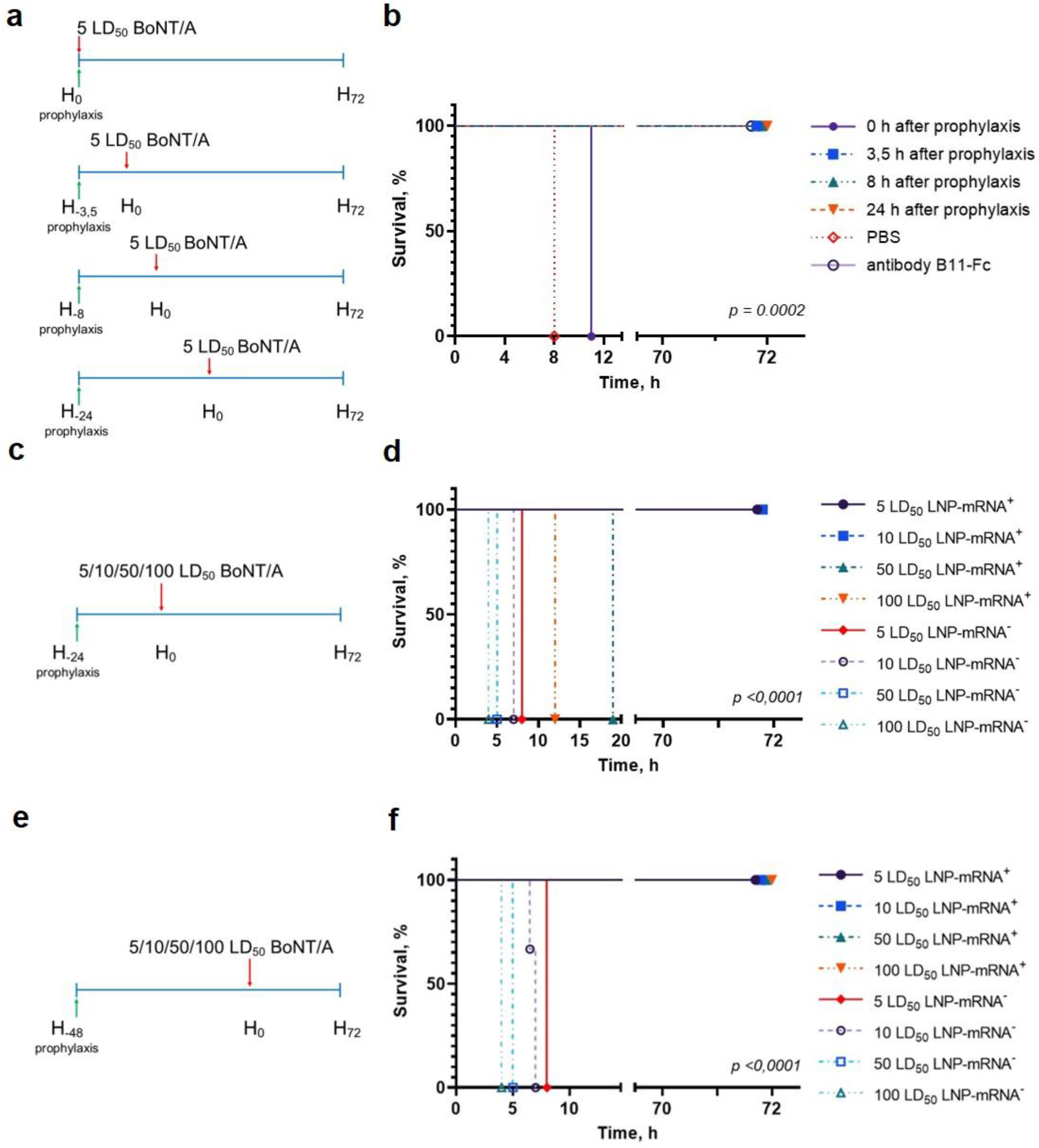
Study of mRNA-B11-LNP prophylactic effect. **a**. Schedule of passive B11-mRNA-LNP immunization before BoNT/A challenge. A total of three mice per group were challenged. Mice received a single intramuscular (i.m.) injection of 10 μg of B11-mRNA-LNP or recombinant B11-Fc antibody 0, 3.5, 8 or 24 h before 5 LD_50_ intraperitoneally injected BoNT/A challenge. Body weight, clinical score and survival were monitored for up to 72 h**; b**. Survival curve of mice that were treated according to the schedule in (**a**). **c**. and **e**. Schedules of passive B11-mRNA-LNP immunization (10 μg of B11-mRNA-LNP per mouse) before challenge by different BoNT/A doses (5, 10, 50 and 100 LD_50_). A total of three mice per group were challenged. Body weight, clinical score and survival were monitored for up to 72 h**; d**. and **f**. Survival curves of mice that were treated according to the schedules in (**c**) and (**e**). A total of three mice per group were challenged. LNP-mRNA^+^ – mice that were treated with B11-mRNA-LNP; LNP-mRNA^−^ – mice that did not receive B11-mRNA-LNP, but were challenged. *p*-value determined by log-rank (Mantel-Cox) test.

Then, we studied the protective effect of B11-mRNA-LNP (injected 24 h and 48 h before challenge) against different BoNT/A doses (5, 10, 50 and 100 LD_50_) according to schedules in **Figure 2c, e**. Mice that were injected with B11-mRNA-LNP 24 hours before challenge did not survive the exposure of BoNT/A high doses (50 and 100 LD_50_) in contrast to those received B11-mRNA-LNP 48 hours before challenge (**Figure 2d, f**).

## DISCUSSION

Recent advances in the treatment and prevention of infectious diseases using mRNA-related approaches have increased the interest in the production of antibodies based on this technology, for both therapeutic and prophylactic purposes. Compared to DNA vectors, the use of mRNA-encoded antibodies brings advantages such as no risk of genome integration, rapid and highly controllable expression, and no dependence on the proliferation status of target cells. In comparison to recombinant mAb technology, mRNA-LNP directs endogenous production of the therapeutic protein with native post-translational modifications and does not require biotech production in cell cultures followed by extensive purification and characterization of the protein product.

In this study we created and applied a new mRNA platform for the production of BoNT/A-specific nanobody fused to the human Fc fragment, and showed its high protective activity in mice. To reduce immunogenicity and provide a prolonged and efficient protein synthesis, in our transcripts we replaced 100% U with m^1^Ψ, and used 5′ cap-1 structure, efficient human *HBA1* 5′ and 3′ UTRs, and a 100-nt long poly(A)-tail, as suggested in earlier studies [24,26–29]. In particular, the use of non-immunogenic mRNA is crucial, because a number of cell innate immune receptors recognizing exogenous RNA can trigger the release of type I interferons leading to activation of interferon inducible genes and inhibition of translation.

The construct was initially characterized in cultured cells (including immortalized human cell lines and primary mouse cells) using luciferase reporter assay. We demonstrated efficient reporter activity during at least 24 h. Notably, the expression was well pronounced only for cap-1 and 100%-m^1^Ψ containing transcripts in the case of dendritic cells, where the innate immune receptors are known to be especially active (**Figure S1**). These results are consistent with previous studies, where mRNA-transfected cells demonstrated a similar dynamics of the Fluc activity [30,31]. However, we would like to note that the continuous real-time measurement of the luciferase activity in living cells we applied [25] provides a more accurate and comprehensive analysis of mRNA expression than the end-point measurements (implying cell lysis) regularly used by others.

Next, we evaluated the effects of the Fluc-mRNA *in vivo* using LNP delivery system. Recent advances demonstrated that LNP provides a wide range of possibilities for researchers in molecular therapeutics. LNP has been specifically developed for delivery into the liver [32] and has previously been shown to be very effective for this purpose, providing robust protein synthesis [31,33]. In this study we used ionizable lipid ALC-0315 combined with DSPC, cholesterol, and a PEG–lipid, a mixture similar to the delivery system used in the Pfizer-BioNTech COVID-19 vaccine [34]. We observed the high level and prolonged expression of Fluc mRNA formulated into LNPs injected through intravenous and intramuscular routes in mice.

As expected, i.v. route allowed obtaining the highest values of luciferase activity in the liver for the entire study period (**Figure 1c, d**). Luminescence was already observed at 15 min after injection and continued about 24 h followed by a decrease. Whereas i.m. injection led to moderate expression, it produced long lasted results: we could detect luminescence for about 3 weeks in the injection site and up to 48 h in the liver (**Figure 1c, e; Figure S2**). Therefore, for our further purposes related to passive immunotherapy, intramuscular administration of mRNA encoding the anti-BoNT/A antibody seemed to be the most appropriate. In general, our data are in good agreement with previously reported results [31] with difference in translation duration, due to higher mRNA-LNP dose used in our study (10 vs. 5 μg per mice).

Next, we analyzed the *in vivo* effects of a similar mRNA construct encoding the B11-Fc antibody. This clone of the camelid sdAb fused with the human Fc fragment was described earlier [20]. Injection of B11-Fc mAb 1 and 3 h in advance or 1 h after 10 LD_50_ BoNT/A administration led to 100% survival [20].

Our results of the experiment with BoNT/A challenge confirmed that mRNA technology can be used for passive immunization against botulism. According our prophylaxis scheme, we have established the time (3.5 h) after B11-mRNA-LNP administration, at which the protective effect was observed though mild symptoms of intoxication began already to show (**Figure 2c**). The presence of clinical manifestations of mild severity indicates that a further reduction in 3.5-hour interval between treatment and challenge may lead to a lethal outcome. Furthermore, we demonstrated that 48-hour mRNA translation ensured the 100% survival even following 100 LD_50_ BoNT/A exposure (**Figure 2f**).

Previously, it has been demonstrated that expression of B11-Fc antibody, delivered via recombinant adeno-associated virus (rAAV-B11-Fc), can protect animals from lethal dose (10 LD) of BoNT/A, starting only from day 3 after administration [11]. Moreover, rAAV-B11-Fc did not ensure survival against higher doses of BoNT/A (50 and 100 LD_50_) [11]. In a recent study, Thran and co-authors showed therapeutic potential of mRNA-mediated antibody expression for animal protection against a number of toxins including BoNT/A [12], challenging mice with 4 LD_50_ of BoNT/A. Our data showed that passive immunization with B11-mRNA-LNP provides rapid (as early as 3.5 h) and robust protective antibody production sufficient to neutralize even significantly higher dose of BoNT/A, which is crucial for emergency prevention.Taken together, these results indicate that mRNA-mediated nanobody expression can be used for both therapeutic and prophylactic purposes in BoNT/A challenge.

### Limitation

In this study, we have not described pharmacokinetics of the target antibody in details, as the currently used test system for measuring antibody levels does not have sufficient sensitivity and specificity. These data should be obtained in future experiments.

### Conclusion

We have developed an mRNA platform for effective gene delivery in mammals. The platform allows long-term expression of mRNA after intramuscular and intravenous administration in mice. Injection of mRNA-LNP encoding the humanized single-domain VHH-Fc antibody specific for botulinum neurotoxin A provides protective activity as early as 3.5 h after administration. Thus, the mRNA-based approach for single-domain antibody delivery paves the way for the development of new immunotherapy drugs for emergency prevention.

## METHODS

### 1. Cloning, the production and purification of plasmid DNA, *in vitro* transcription

The coding regions for the firefly luciferase (from the vector pGL4.51[*luc2*/CMV/Neo] (Promega)) or the camelid anti-BoNT/A sdAb (clone B11) [20] were cloned into a vector for *in vitro* transcription (IVT) based on pJAZZ-OK linear bacterial plasmid **(Figure S3a)**. For this, the *Col*E1 replication origin was first amplified from plasmid pTZ57R (Thermo Fisher Scientific), appended with the *Sma*I restriction site at one end, and inserted into pJAZZ-OK using BigEasy® v2.0 Linear Cloning Kit (Lucigen) to increase plasmid copy number **(Figure S3b)**. The resulting plasmid was then digested with *Sma*I (Sibenzyme) and assembled with PCR product containing mRNA template elements (T7-promoter, the *HBA1* 5′ UTR appended with Kozak sequence [27], the *HBA1* 3′ UTRs, and 100-nt poly(A)-tail followed by BsmBI site) using Gibson Assembly® Ultra kit (Codex) (**Figure S3c**). Before the assembly, SmaI site was inserted between 5′ and 3′ UTRs using Overlap Extension PCR from pT7-5′Luc3′-pA template (unpublished) and BsmBI site in B11-Fc CDS was destroyed using the site-directed mutagenesis resulting in alternative synonymous codon for V193. Finally, after treatment of the resulting plasmid with SmaI, either Fluc or B11-Fc CDSs were cloned directly between 5′ and 3′ UTRs, also using Gibson assembly (**Figure S3d**), with the SmaI site elimination during this process. Cloning steps and plasmid production were performed using freshly prepared *E. coli* BigEasy™-TSA™ Electrocompetent Cells (Lucigen). All cloning procedures were confirmed by Sanger sequencing using BigDye® Terminator v3.1 Cycle Sequencing kit on 3500 Genetic Analyzer (Applied Biosystems). The primer sets used in this study listed in **Table S1**.

The pDNA for IVT were isolated and purified from a culture of the *E. coli* grown for 16 h in 250 ml of 2×YT medium using the Plasmid Maxi Kit (QIAGEN) in accordance with the manufacturer’s protocol. The pDNA digestion was performed using BsmBI-v2 restriction endonuclease (NEB), followed by purification of the linearized plasmid by phenol-chloroform extraction and ethanol precipitation. IVT was performed as described earlier [21], with the following modifications. 100-μl reaction volume contained 3 μg of DNA template, 3 μl T7 RNA polymerase (Biolabmix) and 10×Buffer (TriLink), 4 mM trinucleotide cap 1 analog ((3′-OMe-m^7^G)-5′-ppp-5′-(2′-OMeA)pG)) (Biolabmix), 5 mM m^1^ΨTP (Biolabmix) replacing UTP, and 5 mM GTP, ATP and CTP. After 2 h incubation at 37 °C, 6 μl DNase I (Thermo Fisher Scientific) was added for additional 15 min, followed by mRNA precipitation with 2M LiCl (incubation for 1 h in ice and centrifugation for 10 min at 14,000 g, 4 °C) and carefully washed with 80% ethanol. RNA integrity was assessed by electrophoresis in 8% denaturing PAGE.

### 2. Formulation of mRNA in LNP

*In vitro* transcribed mRNAs were encapsulated in LNPs using a self-assembly process in which an aqueous solution of mRNA at pH 4.0 is rapidly mixed with a solution of lipids dissolved in ethanol. LNPs used in this study were similar in composition to those described previously [18,31,34]. Briefly, lipids were dissolved in ethanol at molar ratios of 46.3:9:42.7:1.6 (ionizable lipid:distearoyl PC:cholesterol:PEG-lipid) [34]. Acuitas ionizable lipid (ALC-0315) and PEG-lipid (1,2-Dimyristoyl-sn-glycero-3-methoxypolyethylene glycol 2000) were purchased in Cayman Chemical Company. The lipid mixture was combined with an acidification buffer of 10 mM sodium citrate (pH 4.0) containing mRNA (0.2 mg/ml) at a volume ratio of 3:1 (aqueous: ethanol) using a Nanoassemblr Spark device (Precision NanoSystems). The ratio of ionizable nitrogen atoms in the ionizable lipid to the number of phosphate groups in the mRNA (N:P ratio) was set to 6 for each formulation. Formulations were dialyzed against PBS (pH 7.2) in Slide-A-Lyzer dialysis cassettes (Thermo Fisher Scientific) for at least 24 h. Formulations were concentrated using Amicon ultra-centrifugal filters (EMD Millipore), if needed, then passed through a 0.22-μm filter and stored at 4 °C (PBS) until use. Formulations were tested for particle size, zeta potential, RNA encapsulation. All LNPs were found to be about 80 nm in size as measured by dynamic light scattering using a Zetasizer Nano ZS instrument (Malvern Panalytical). mRNA encapsulation was about 90% (measured by RiboGreen assay – Quant-iT™ RiboGreen™ RNA Reagent, Thermo Fisher Scientific).

### 3. Cell culture, transfection and luciferase measurements

HeLa and HEK293T cells were obtained from ATCC and cultured in DMEM (Gibco) at 37 °C, in a 5% CO_2_ humidified atmosphere. Bone marrow derived dendritic cells (BMDCs) from C57BL/6 mice were differentiated from proliferating mouse bone marrow progenitors through induction with 20 ng/ml granulocyte macrophage colony stimulating factor (GM-CSF) (R&D Systems, USA) over 6–9 days as described [35]. Briefly, mice were euthanized by CO_2_ overdose and the femurs and tibias were collected in ice-cold Hank’s balanced salt solution (HBSS, Sigma-Aldrich). The muscles were removed with a scalpel and by rubbing the bones with a tissue. The ends of the bones were then cut off with scissors and crushed. The bone marrow was flushed out with 2–3 ml of RPMI complete medium in a syringe with a 25-gauge needle. All bone marrow cells were collected and washed twice with HBSS. The bone marrow cells were cultured in 24-well plates containing approximately 5×10^5^ cells/ml in 1 ml total volume. The cells were maintained at 37 °C with 5% CO_2_ in complete RPMI medium with 10% heat inactivated fetal calf serum (PAA), 0.05 mM 2-mercaptoethanol (Thermo Fisher Scientific), 1×Non Essential Amino Acids (PanEco), 20 ng/ml GM-CSF, 2 mM glutamine, 100 U/ml penicillin, and 100 μg/ml streptomycin (all PanEco). After 24 hours, the nonadherent cells were collected and discarded, and 1 ml of fresh media was added to each well. On day 3, 5 and 7 half of the medium in each well was replaced with fresh medium. On day 7–9, the percentage of CD11c-positive cells in non-adherent population in the cultures was ∼60–70%. Mouse primary fibroblasts (MPF) were derived from auricles according to the protocol [36]. Briefly, the animals were euthanized, their auricles were collected and rinsed with 70% ethanol. Tissue fragments were dissected with sterile scissors in a FBS-free DMEM solution containing 1x Liberase TM Research Grade (Roche) and Antibiotic-Antimycotic (Gibco), and incubated at 37 °C on a shaker for 1 h. The tissue fragments were then centrifuged thrice in 50 ml of growth media (DMEM/F12 with 1×Antibiotic-Antimycotic and 15% FBS (Gibco) for 5 min at 500 g, the pellet was resuspended in 10 ml of growth media and incubated for 7 days in a tissue culture dish at 37 °C, 5% CO_2._ After 7 days the cells were trypsinized, plated in a new culture dish, and used for experiments within the first 12 population doublings.

Transfection with the Fluc encoding mRNA was performed essentially as described previously [25]. Shortly, a day before transfection, ∼3×10^4^ cells per well were transferred into the white FB/HB 96-well plates (Greiner) in 75 μl of DMEM (Paneco) supplemented with 10% FBS (HyClone) in the presence of penicillin and streptomycin (Paneco). Next day, cell culture at ∼70% confluency was transfected with the reporter mRNA. For this, 30 ng of mRNA in 20 μl Opti-MEM (Gibco) per well was mixed with a solution of 0.06 μl of GenJector™-U (Molecta) in 3 μl Opti-MEM (per well), incubated for 15 min, then supplied with 0.4 μl of 100 mM D-luciferin (Promega) per well, and then 23 μl of the mixture were added to the cells. During these steps, all manipulations were performed as described in the FLERT protocol [22], providing a rapid and non-stressful transfection procedure. Real-time luminescence measurements were carried out overnight in the CLARIOstar plate reader (BMG Labtech) equipped with Atmospheric Control Unit to maintain 5% CO_2_, at 37 °C (signal integration time – 5 s). All the experiments were repeated at least three times (including ones with different cell passages), the mean values ± SD were calculated.

### 4. Animal experiments

All animal experiments were performed in accordance to the Directive 2010/63/EU (Directive 2010/63/EU of the European Parliament and of the Council of 22 September 2010 on the Protection of Animals Used for Scientific Purposes. Off J Eur Communities L276:33–79), FELASA recommendations [37], and the Inter-State Standard of «GLP» (GOST - 33044-2014) approved by the Institutional Animal Care and Use Committee (IACUC) of the Federal Research Centre of Epidemiology and Microbiology named after Honorary Academician N.F. Gamaleya and were performed under Protocol #22 from 15 April 2022. All persons using or caring for animals in research underwent training annually as required by the IACUC.

Females BALB/c mice, 5 – 6-week-old, weighting 18 – 20 g were randomly allocated into groups. Fluc-mRNA LNPs in PBS, or PBS (for placebo group) were injected by two ways of administration, intravenous and intramuscular, in dose 10 μg mRNA per mouse. Following 30 min, 3 and 24 h after Fluc mRNA LNPs injection, 2.5 μg of D-luciferin was injected intraperitoneally. For intramuscular administration, bioluminescence was measured for 24 days (additional points – 2, 3, 4, 7, 10, 14, 21, 24 days). Then, after 5 min exposition, bioluminescence in mice was visualized using IVIS Lumina II (Life sciences) under isoflurane anesthesia. Animals (n = 3 per group) that were not used for *in vivo* visualization, were euthanized by CO_2_ inhalation with subsequent isolation of internal organs. The firefly luciferase activity was detected in homogenates of organs by luminescence measurements using Synergy H4 hybrid reader (Bio-Tek) and Bright-Glo Luciferase Kit (Promega).

### 5. Botulinum Toxin A preparation

Botulinum toxin A was obtained using *C. botulinum* strain A98. The strain was cultivated under anaerobic conditions for five days. Bacterial suspension was precipitated by centrifugation at 5000×g for 30 min at 10 °C. Bacterial proteins from the filtrate were concentrated by trichloroacetic acid precipitation at pH = 3.8 for 45 min. The precipitate was separated by centrifugation at 12000×g for 30 min at 10 °C and dissolved in 47 mM citrate-phosphate buffer with pH = 5.6. S300 gel filtration and ion-exchange chromatography were performed on the AKTA start system (GE Healthcare Life Science) with DE cellulose (Pharmacia). The toxin (150 kDa) was additionally purified by chromatography with DE cellulose (Pharmacia) in borate buffer with pH = 8 and eluted by 50 mM NaCl. The specific antigenic activity of BoNT/A was determined by reaction with monospecific antibodies to BoNT/A (Gamaleya Laboratory of Clostridiosis and commercial preparation of Scientific Centre for Expert Evaluation of Medicinal Products Russian Federation). Purified BoNT/A was checked by SDS-electrophoresis with 2-mercaptoethanol. The toxin was filtered through 0.22 μm syringe filter before injection. The BoNT/A concentration was determined spectrophotometrically (UV 280 nm) and calorimetrically (Bradford protein analysis). LD_50_ was determined on six-week-old female BALB/c mice (weighing 18 – 20 g). Specific activity was determined by the standard mouse lethality assay described by Lindstrom and Korkeala [38]. The specific activity ranged from 10 – 30 pg/mouse among lots of BoNT/A. Variations in the specific activity of BoNT/A was associated with different batches of animals and BoNT/A. Therefore, we checked the specific toxic activity before each experiment and for each batch of animals and the toxin.

### 6. Challenge experiments

Five-to six-week-old BALB/c mice weighting 18 – 20 g were randomly allocated into groups (three animals per group). Mice from all groups were housed in standard isolator cages under standard conditions and were challenged with 5 LD_50_ (or 10, 50, 100 LD_50_) dose of intraperitoneal injection of BoNT/A. For prophylactic treatment a single intramuscular injection of LNP-formulated B11-mRNA, or PBS (for negative control group), or recombinant B11-Fc antibody [20] in a dose of 0.5 mg/kg (for positive control group) were administered 0 / 3.5 / 8 / 24 h before BoNT/A challenge. For high dose survival study B11-mRNA-LNP was injected 24 and 48 h before challenge. Mice were monitored for clinical symptoms at intervals: 0, 4, 8, 26, 48 and 72 h after challenge. Assessment of clinical symptoms was carried out in accordance with the scale: score of 0 (no symptoms), score of 1 (mild symptoms), score of 2 (moderate symptoms), score of 3 (severe symptoms = humane endpoint). Time of death was defined as the time at which a mouse was found dead or was euthanized via carbon dioxide asphyxiation followed by cervical dislocation at humane endpoint.

### 6. Statistical analysis

Data were analyzed using GraphPad Prism software version 8.

## ACKNOWLEDGEMENTS

We are grateful to Timofey A. Remizov and Anastasia A. Zakharova for their help with the purchase of reagents and paperwork for ongoing projects. We thank Dr. Anatoly N. Noskov for his valuable advice and materials necessary for setting up an BoNT/A challenge animal model. We are also grateful to Maria Konopleva for assistance with cell culture. E.A.P., I.I.S., and S.E.D. are members of the Interdisciplinary Scientific and Educational School of Moscow University “Molecular Technologies of Living Systems and Synthetic Biology”.

## SUPPLEMENTARY MATERIAL

**Figure S1.**
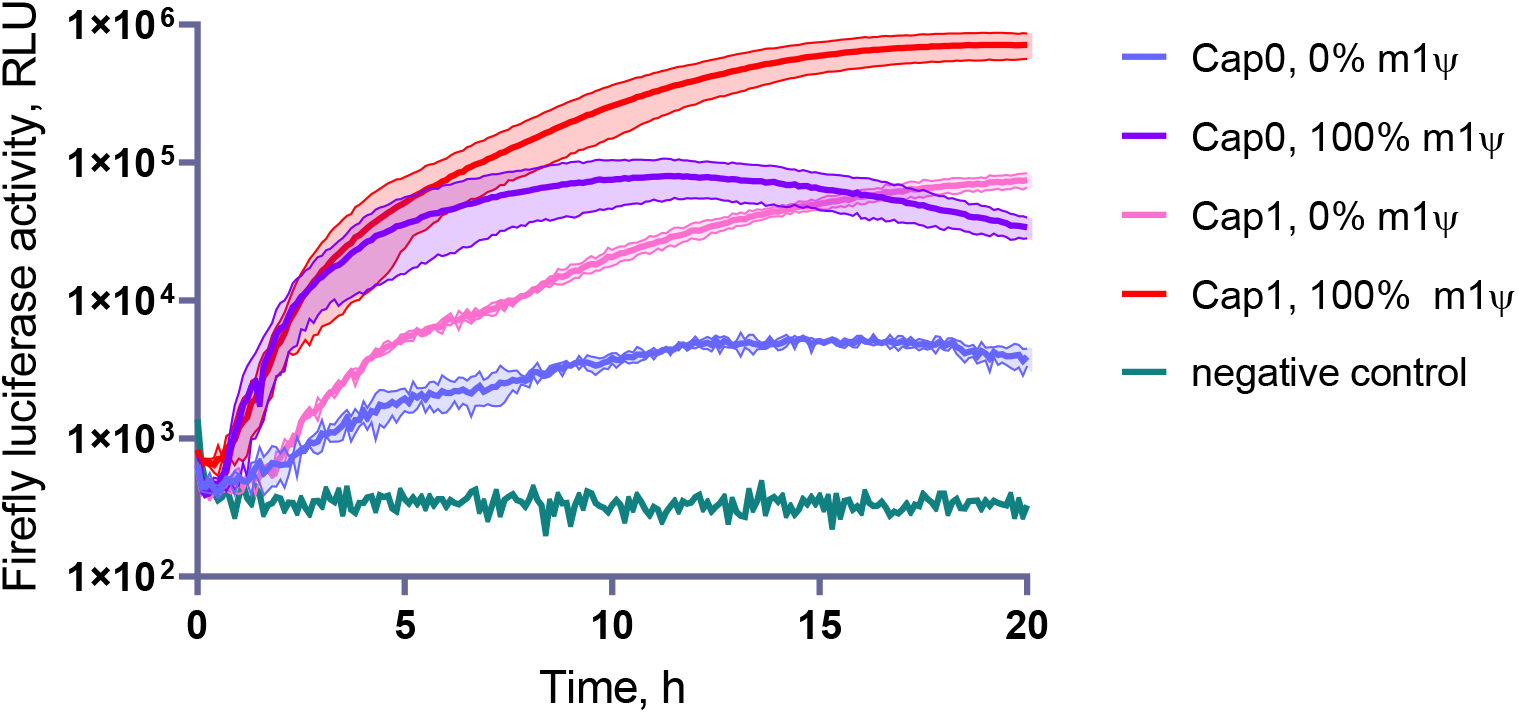
The influence of the presence of a cap1/cap0 structure and incorporation of modified nucleoside, N1-methylpseudouridine (m^1^Ψ), on the firefly luciferase mRNA expression in mouse bone marrow-derived dendritic cells (mBMDC).

**Figure S2.**
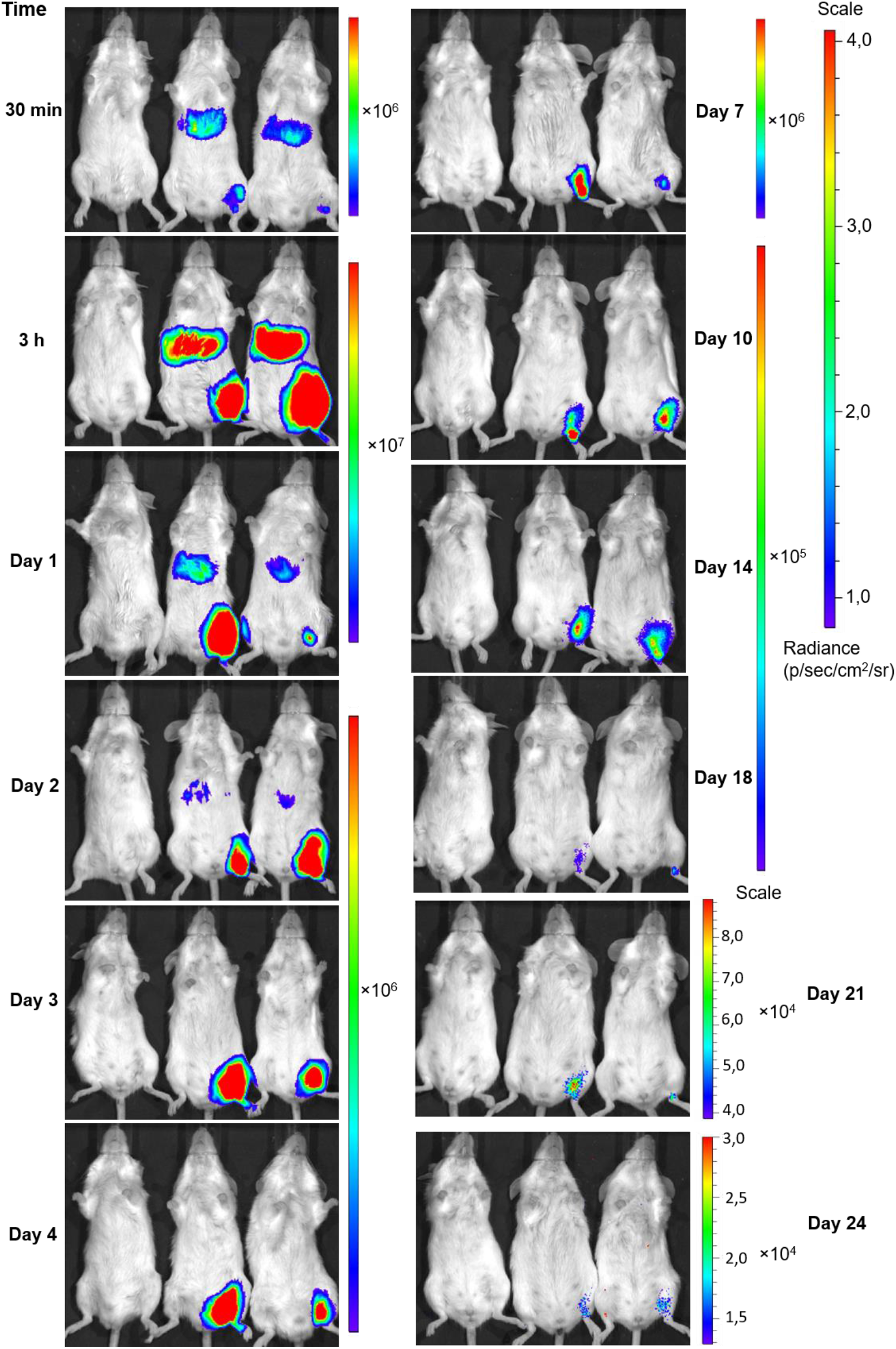
Representative IVIS images of BALB/c mice injected with 5.0 μg mRNA-LNP in PBS, or PBS (for placebo group) by intramuscular (i.m.) route. Relative luminescence plot is shown, and the scale of luminescence is indicated.

**Figure S3.**
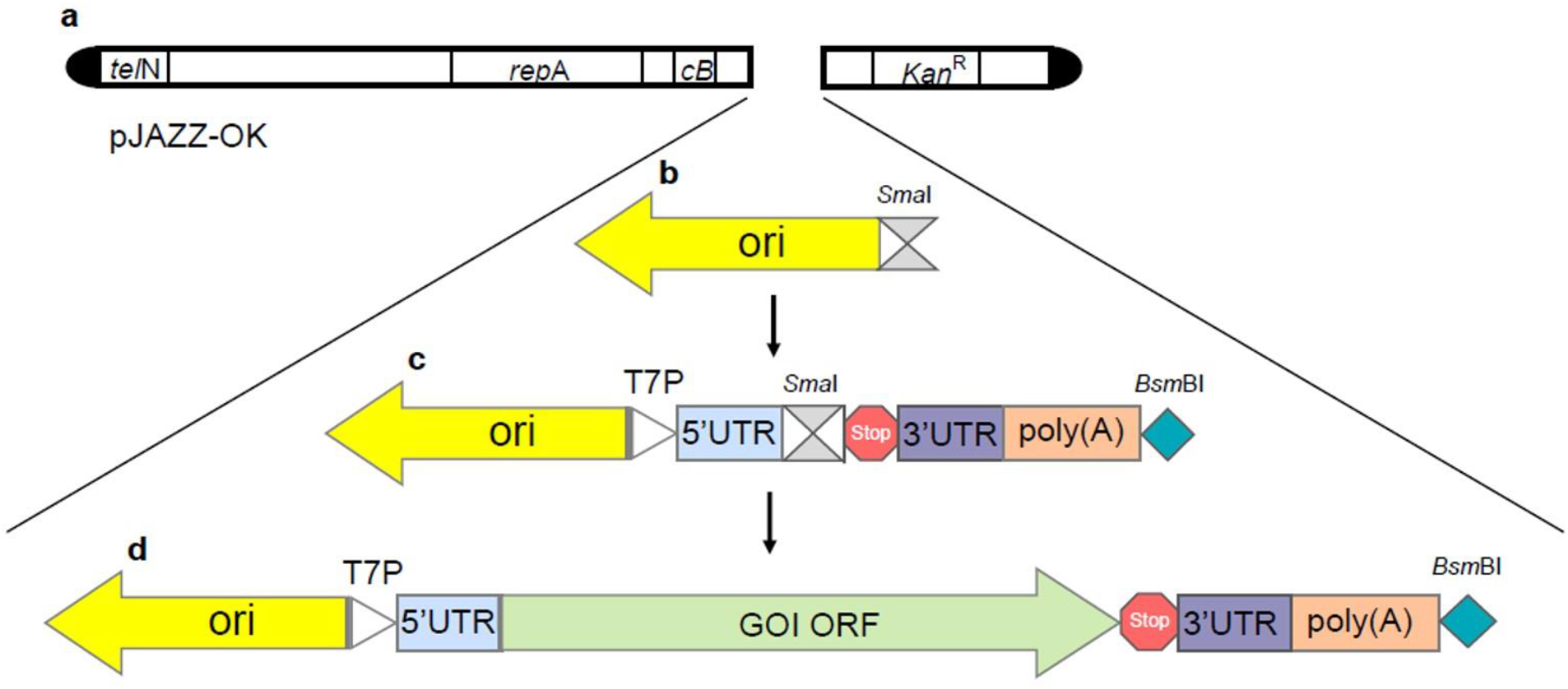
Cloning strategy for IVT plasmid templates based on pJAZZ-OK. (Details described in Methods section 1). **a**. Schematic of commercial pJAZZ-OK linear bacterial plasmid derived from N15 bacteriophage genome (*telN* – protelomerase gene, *rep*A – replication factor gene and origin of replication, *cB* – the primary replication repressor protein, *Kan*^R^ – kanamycin resistance gene, black semi-circles denoting terminal hairpin loops); **b**. Schematic of linear plasmid with *Col*E1 origin of replication (*ori*, yellow arrow) followed by *Sma*I restriction site (grey block); **c**. Schematic of plasmid contained backbone sequences for IVT: T7-promoter (*T7P*, white triangle), human *HBA1* 5′ and 3′ UTRs (blue and plum blocks, respectively) cleaved by *Sma*I site, 2x strong translation terminator (*Stop*, red octagon), 100-nt poly(A)-tail (orange block) followed by *BsmB*I restriction site (cyan rhombus); **d**. Schematic of resulted linear plasmid with open reading frame of luciferase or B11 antibody (*GOI ORF*, green arrow).

**Table S1.**
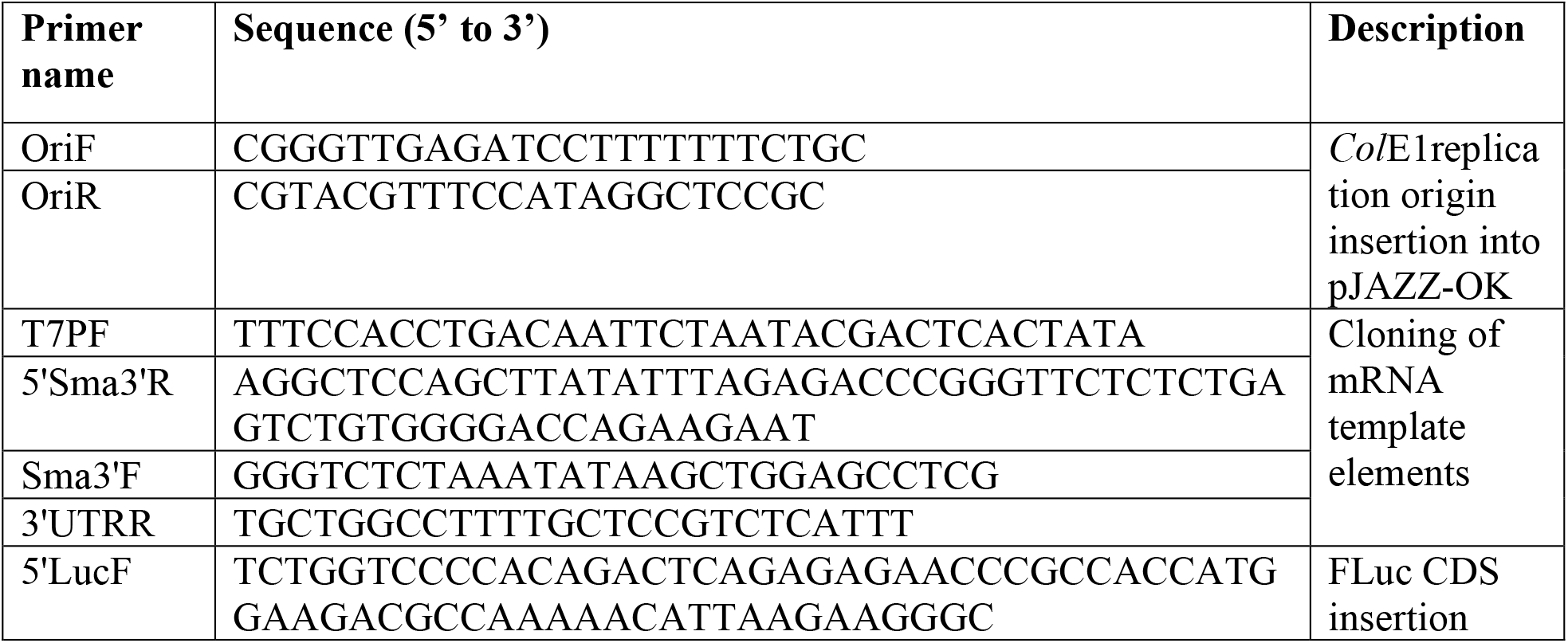
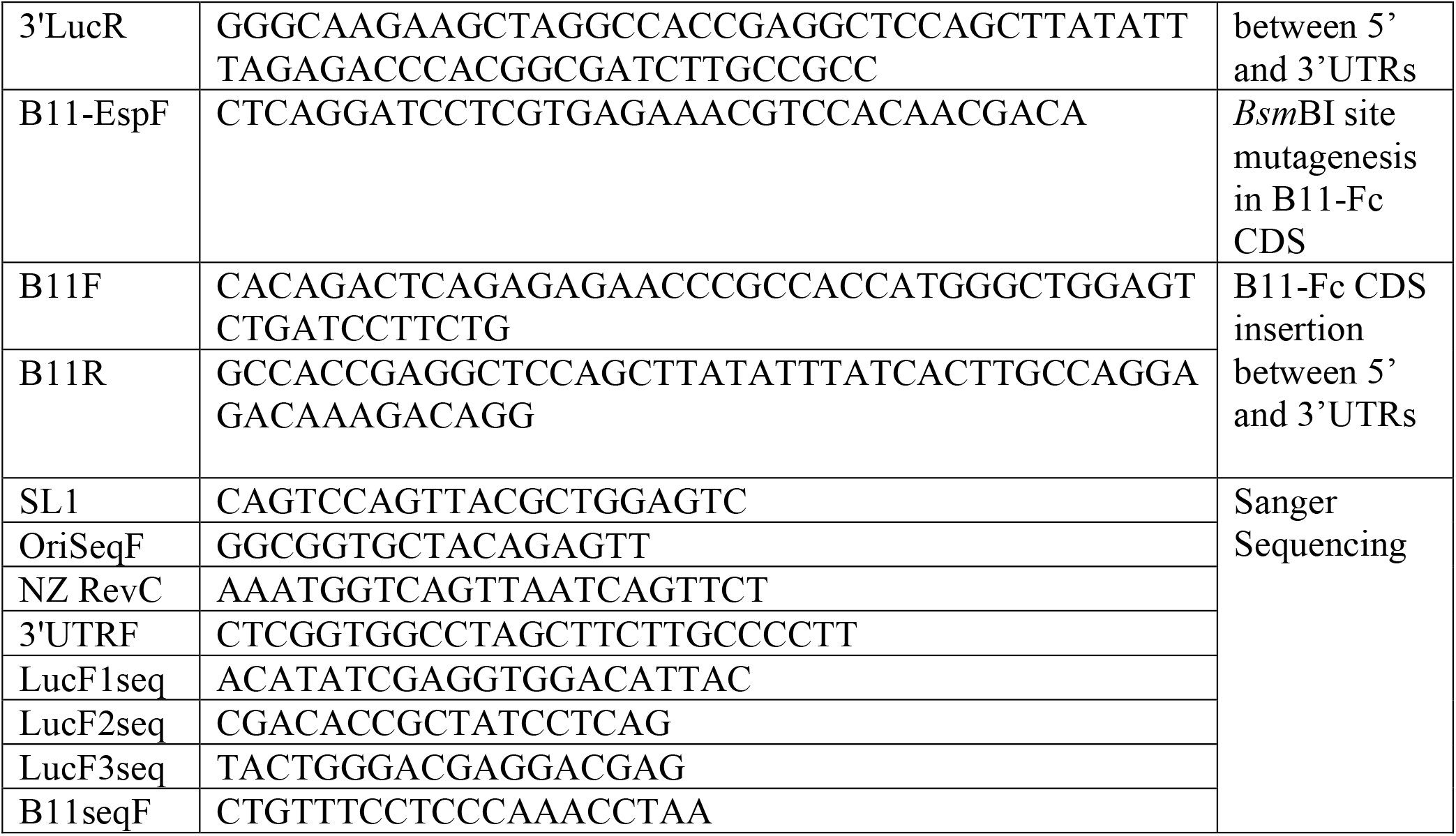
The primer sets used in the study.

